# Developmental and environmental plasticity in opsin gene expression in Lake Victoria cichlid fish

**DOI:** 10.1101/2021.09.01.458542

**Authors:** Lucia Irazábal-González, Daniel Shane Wright, Martine Maan

**Affiliations:** Groningen Institute for Evolutionary Life Sciences (GELIFES), University of Groningen, Groningen, The Netherlands; Division of Evolutionary Biology, Ludwig Maximilian University of Munich, Munich, Germany

**Author notes:** Corresponding author: Lucía Irazábal-González, Groningen Institute for Evolutionary, Life Sciences, University of Groningen, Groningen, The Netherlands.

**Keywords:** *Pundamilia*, vision, adaptation, light, heterochrony

## Abstract

In many organisms, sensory abilities develop and evolve according to the changing demands of navigating, foraging and communication across different environments and life stages. Teleost fish inhabit heterogeneous light environments and exhibit a large diversity in visual system properties among species. Cichlids are a classic example of this diversity, generated by different tuning mechanisms that involve both genetic factors and phenotypic plasticity. Here, we document the developmental progression of visual pigment gene expression in Lake Victoria cichlids and test if these patterns are influenced by variation in light conditions. We reared two sister species of *Pundamilia* to adulthood in two distinct visual conditions that resemble the two light environments that they naturally inhabit in Lake Victoria. We also included interspecific first-generation hybrids. We then quantified (using RT-qPCR) the expression of the four *Pundamilia* opsins (SWS2B, SWS2A, RH2A and LWS) at 14 time points. We find that opsin expression profiles progress from shorter-wavelength sensitive opsins to longer-wavelength sensitive opsins with increasing age, in both species and their hybrids. The developmental trajectories of opsin expression also responded plastically to the visual conditions. Finally, we found subtle differences between reciprocal hybrids, possibly indicating parental effects and warranting further investigation. Developmental and environmental plasticity in opsin expression may provide an important stepping stone in the evolution of cichlid visual system diversity.

**Research highlights:** In Lake Victoria cichlid fish, expression levels of opsin genes (encoding visual pigments) differ between developmental stages and between experimental light treatments. This plasticity may contribute to the evolution of cichlid visual system diversity.

## Introduction

Animal sensory systems mediate interactions with the environment, contributing to foraging, navigation, predator avoidance and mate selection. Sensory systems are highly diverse within and between species, associated with differences in ecological niche and life history (Stevens, 2013). For example, the visual system rapidly adapts to variation in local light conditions, resulting in inter- and intraspecific variation in visual system properties (Bowmaker, 2008; Carleton, Dalton, Escobar[Camacho, & Nandamuri, 2016; Chang et al., 2021; Veilleux & Kirk, 2014). In water, where light transmission is poorer than on land, there is substantial heterogeneity in light conditions, mediated by water depth and transparency. As a result, variation in visual properties is high across aquatic vertebrates, compared to other groups (Bowmaker et al., 1994; Loew & McFarland, 1990).

Cichlids are a family of teleost fish that has rapidly diverged into numerous species across Africa, Asia, South America and Central America (Kocher, 2004; Salzburger, 2018; Seehausen, 2006). The species inhabit different photic environments, and their visual system properties are highly diverse (Carleton & Yourick, 2020). Some of this diversity is genetically encoded, such as variation in the opsin gene sequences that affect the wavelength absorption properties of the visual pigments, and the subset of opsin genes actually expressed (Carleton et al., 2016; Carleton, Parry, Bowmaker, Hunt, & Seehausen, 2005; Carleton et al., 2008; Hofmann et al., 2009). Other aspects of the visual system can be influenced by developmental or environmental plasticity, including the expression levels of the opsin genes and the use of alternative chromophores (Carleton et al., 2016; Carleton & Kocher, 2001; Halstenberg et al., 2005; Härer, Torres[Dowdall, & Meyer, 2017; Hofmann et al., 2009). In this study, we examine the developmental trajectory of relative opsin expression in two Lake Victoria cichlid species (genus *Pundamilia*) and their hybrids and investigate the influence of light-induced plastic variation in opsin expression.

Cichlids have seven distinct cone opsin genes: the short-wavelength-sensitive opsins SWS1 (UV), SWS2B (violet), SWS2A (blue); the medium-wavelength-sensitive opsins RH2Aα, RH2Aβ, RH2B (green); and the long-wavelength-sensitive opsin LWS (red) (Carleton et al., 2016; Carleton et al., 2005). During development, opsin expression profiles may change, and species differ in their developmental patterns. This variation (heterochrony) may provide a target for divergent selection and contribute to the evolution of differences in opsin gene expression observed across species (Carleton et al., 2008; Härer et al., 2017; O’Quin, Smith, Sharma, & Carleton, 2011; Sandkam et al., 2020).

Carleton et al. (2008) characterised the developmental patterns of opsin expression in two Lake Malawi cichlid clades (sand-dwelling *Dimidiochromis compressiceps* and *Tramitichromis intermedius* and rock-dwelling *Metriaclima zebra ‘gold’, Metriaclima zebra, Metriaclima benetos*, and *Labeotropheus fuelleborni*) and an ancestral riverine cichlid lineage (Tilapia, *Oreochromis niloticus*). *O. niloticus* expressed predominantly short-wavelength-sensitive opsins as larvae (SWS1 and RH2B), increased the expression of medium-wavelength-sensitive opsins as juveniles (SWS2B and RH2A) and finally expressed high amounts of long-wavelength-sensitive opsins as adults (SWS2A, RH2A and LWS) (Carleton et al., 2008). Rock-dwelling Lake Malawi larvae and juveniles expressed short-and medium-wavelength-sensitive opsins (SWS1, SWS2B and RH2A) and maintained this expression pattern until adulthood. Larvae and juveniles of the sand-dwelling species expressed long-wavelength-sensitive opsins (SWS2A, RH2A and LWS) and maintained this expression profile throughout development. Carleton et al. (2008) hypothesised that the developmental pattern of opsin expression in Lake Victoria cichlids resembles that observed in the Lake Malawi sand-dwellers, but the available data comes from only a small number of Lake Victoria juveniles sampled at a single time point (Carleton et al., 2005; Carleton et al., 2008). Therefore, the developmental pattern of opsin expression in Lake Victoria cichlids remains to be established.

*Pundamilia pundamilia* and *Pundamilia nyererei* are two closely related cichlid species that co-occur at several rocky islands in southeast Lake Victoria, in the Mwanza and Speke Gulfs (Seehausen et al., 2008). Water transparency differs between sites, with turbid waters in the south and clearer waters in the north (Castillo Cajas, Selz, Ripmeester, Seehausen, & Maan, 2012). Both species are consistently depth-segregated at all locations but with more overlap in more turbid locations. *P. pundamilia* is restricted to shallow waters (0 - 2 m), while *P. nyererei* is most abundant at 3 - 10 m depth, where the light spectrum is shifted towards longer wavelengths and is largely lacking short-wavelength light (Seehausen et al., 2008). Anatomically, both species are similar, but they differ in male coloration. *P. pundamilia* males are metallic blue with a blue dorsal fin, and *P. nyererei* males are red with yellowish flanks. Females of both species are cryptic yellow/grey.

In this study, we use fish from Python Island, where until recently, the blue and red species were thought to be *P. pundamilia* and *P. nyererei*, respectively. However, Meier et al. (2017; 2018) showed that the population at Python Island represents a separate speciation event and is, therefore, referred to as *P*. sp. ‘*pundamilia-like*’ and *P*. sp. *‘nyererei-like’*. Across *Pundamilia* populations, and consistently linked to male colour, depth, and water clarity, the LWS opsin gene has sequence variations that alter pigment sensitivity (Hofmann et al., 2009; Seehausen et al., 2008; Wright et al., 2019). Wild populations also differ in opsin expression within and between locations (Hofmann et al., 2009; Wright et al., 2019). In the laboratory, we mimicked the natural shallow (broad-spectrum) and deep (red-shifted) light environments of Python Island and observed evidence of opsin expression plasticity: adult fish expressed more LWS and less SWS2A when reared in red-shifted light conditions, compared to their counterparts from broad spectrum light conditions (Wright et al., 2020).

Here, we characterise the developmental patterns of opsin expression in *P*. sp. *‘pundamilia-like’*, *P*. sp. *‘nyererei-like’* and their hybrids. We hypothesise that the developmental patterns of both species will resemble those of the Lake Malawi sand-dwellers and that the hybrids will be intermediate. We also explore the extent of environmental plasticity during development by rearing the fish in broad-spectrum and red-shifted light conditions (Maan, Seehausen, & Groothuis, 2017; Wright et al., 2017). We expect longer wavelength sensitivities in the red-shifted light condition compared to the broad-spectrum light condition. Moreover, a study in neotropical Midas cichlids suggests that the transition to the adult expression profile may be delayed in short-wavelength-rich light environments and accelerated in long-wavelength-rich environments (Härer et al., 2017), Therefore, we predict that LWS expression increases faster in the red-shifted light condition. For both overall opsin expression and its developmental pattern, we expect hybrids to respond more strongly to the light treatments than the parental species. This is because they are expected to be heterozygous for species-specific alleles influencing opsin expression, which may allow for greater environmental effects on the phenotype.

## Material and methods

### Fish species and breeding

For this experiment, we used first (F1) and second (F2) generation aquarium-reared offspring of wild caught *P*. sp. *‘pundamilia-like’* and *P*. sp. *‘nyererei-like’* collected from Python Island (−2.6238, 32.8567) in 2014.

For breeding, we housed several females with a single male. All fish were tagged (PIT tags, Passive Integrated Transponder, Biomark, ID, Idaho; USA, and Dorset Identification, Aalten, Netherlands). Haplochromine cichlids are maternal mouthbrooders; about five days after fertilization, the eggs were removed from the mother’s buccal cavity and divided equally between light treatments (described below).

We reared both species, as well as their reciprocal hybrids, in both light environments. The analysed samples came from 33 aquarium-reared F1 and F2 families, with 24 dams and 16 sires (Table S1). The animal experiments that we performed for this study were approved by the animal ethics committee of the University of Groningen (DEC 6205B; AVD105002016464).

### Housing and light conditions

All fish were maintained at 25 ± 1 °C, on a 12:12h day:night light cycle. In both light treatments (mimicking the shallow and deep waters at Python Island), the tanks were lit with halogen bulbs (Philips Halogen Masterline ES, 35W) that were differently filtered depending on the treatment. In the deep treatment, the light was filtered with yellow (no. 15 by LEE, Andover, UK) and green filters (LEE no. 243), generating a red-shifted light spectrum (Fig. S1). In the shallow treatment, the light was filtered with the green filter only. Halogen bulbs have a limited short-wavelength radiance, so the short-wavelengths were supplemented with blue lights (Paulmann 88090 ESL Blue spiral 15W) in the broad-spectrum light environment (Fig. S1). Further details on the light treatments and a comparison to the spectral conditions at Python Island are provided in the supplementary information.

### Samples

Fish were sampled at 14 different time points, ranging from 10 to 180 days post fertilization (dpf) (Table S2; *Pundamilia* reach sexual maturity at ~8 months of age; i.e., ~240 dpf). To control for variation between families, we aimed to include at least two different families at each point. Based on previous studies (Carleton et al., 2008; O’Quin et al., 2011) and on pilot trials of total RNA isolation from fish of various ages, the number of eyes sampled differed between time points: we pooled both eyes from 2 individuals at 10 - 20 dpf, both eyes from 1 individual at 30 - 60 dpf, and used 1 eye from a single fish from 70 dpf onwards. Whole eyes were used up to 90 dpf; from 120 dpf onwards we used retinal tissue.

Fish were euthanized using an overdose of MS-222 buffered with sodium bicarbonate, and the eyes were removed, preserved in RNALater (Ambion®, Foster City, CA, USA) and frozen until qPCR analysis (detailed below). Previous studies have shown a circadian rhythm in opsin expression (Halstenberg et al., 2005). To minimize variation and maximize yield, all samples were collected between 16:00 and 18:00.

### Quantification of opsin expression by RT-qPCR

Following previous studies (Wright et al., 2019; Wright et al., 2020), we isolated total RNA with Trizol (Ambion) and reverse-transcribed one microgram of RNA from each sample with Oligo (dT)_18_ (100μM) (Thermo Scientific, Life Technologies) and RevertAid H minus RT reverse transcriptase (Thermo Scientific, Life Technologies) at 45 °C.

We performed real time quantitative PCR (RT-qPCR) to quantify the relative expression of the four opsins present in adult Lake Victoria cichlids (SWS2B, SWS2A, RH2A and LWS (Hofmann et al., 2009)), using gene-specific TaqMan® primers and probes (Table S2). RH2Aα and RH2Aβ were analysed together as RH2A because they are highly similar in sequence and function (Parry et al., 2005). Fluorescence was monitored during 50 cycles with a BIO-RAD C1000 Thermal Cycler (CFX96 Real Time System) (95 °C for 2 minutes, 95 °C for 50 seconds and 60 °C for 1 minute).

We used LinRegPCR® to determine the critical threshold for each sample. This program examines the log-linear part of the PCR curve for each sample to determine the upper and lower limit of the “window of linearity”. Linear regression analysis is then used to calculate the intercept (i.e., the estimated initial concentration) (Ramakers, Ruijter, Deprez, & Moorman, 2003). We used the same approach to calculate the slope and the intercept of a serially diluted construct of the four opsin genes ligated together. We used the following equation to calculate the relative opsin gene expression:

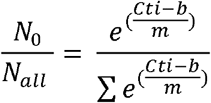

where *N_0_/N_all_* represents the expression of each opsin gene, relative to total opsin expression. *Cti* is the threshold cycle number for the focal opsin, *b* is the intercept of the mean *Cti* of each diluted point of the ligated standard, and *m* the slope of this ligated standard. All samples were run twice.

### Data analysis

We discarded samples with PCR efficiencies below 1.75 and above 2.25 and when the *Cq* standard deviation between replicates was greater than 0.5. To avoid unwarranted exclusion of low expression levels, we did not apply these rules for opsins with < 1% of the total expression (efficiencies decrease and error rates increase at such low concentrations). The relative expression of each opsin is defined in relation to the other three opsins, so discarding a sample for one opsin means discarding the entire sample. The RT-qPCR from the discarded samples was repeated once for any opsin that fell outside of the parameters mentioned above. After applying these quality thresholds, 26 of 239 samples were discarded.

Prior to statistical analysis, we performed an outlier check with Tukey’s method identifying the outliers that fell above and below the 1.5 interquartile range. For this procedure, we divided the dataset by species and treatment. Moreover, due to the change in opsin expression with developmental time (see results), the data was divided by age of sampling (10 to 30 dpf and 40 dpf onwards). This analysis was performed to ensure that the dataset did not contain artefacts from the sampling, RT-qPCR or the methodology (as discussed in Carleton et al., 2005; Hofmann et al., 2009; Wright et al., 2019; Wright et al., 2020). Thirty-four outliers were removed, leaving 179 samples for the analyses. We used linear mixed modelling (in R, R Development Core Team 2017, lmer, package lme4), separately for each opsin, with the time (dpf), species (*P*. sp. *‘pundamilia-like’*, *P*. sp. *‘nyererei-like’* or hybrid) and light treatment (broad-spectrum or red-shifted) as fixed effects, and mother and father ID as random effects (*expression~treatment*species*time+ (1|momID) + (1|dadID)*). We used stepwise backward selection based on statistical significance (in R, ‘MASS’ package version 7.3-45; (Venables & Ripley, 2002)) to determine the minimum adequate model (retaining parental identities in all models to account for pseudo replication). Finally, we performed “KRmodcomp” to estimate the parameter effects, P-values, and degrees of freedom based on the Kenward-Roger approximation (in R, package ‘pbkrtest’ version 0.4-6; Halekoh and Højsgaard (2014)).

## Results

### *Pundamilia* opsin expression throughout development

The expression of all four opsins changed significantly over time, as evidenced by the significant effect of time in all models - particularly during early development (10 - 90 dpf; Fig. 1; Table 1). Throughout development, *Pundamilia* expressed high proportions of LWS, followed by RH2A, SWS2A and SWS2B. With time, RH2A expression decreased from ~25% to below 20%, and LWS increased from ~60% to ~75%. The expression of both SWS2 opsins was low in early development (~5%). SWS2A increased to about 20%, while SWS2B decreased to nearly 0 at 50 dpf. The expression of all four opsins stabilized after 200 dpf, reaching the levels previously established for adult fish (Fig. 1).

**Figure 1.**
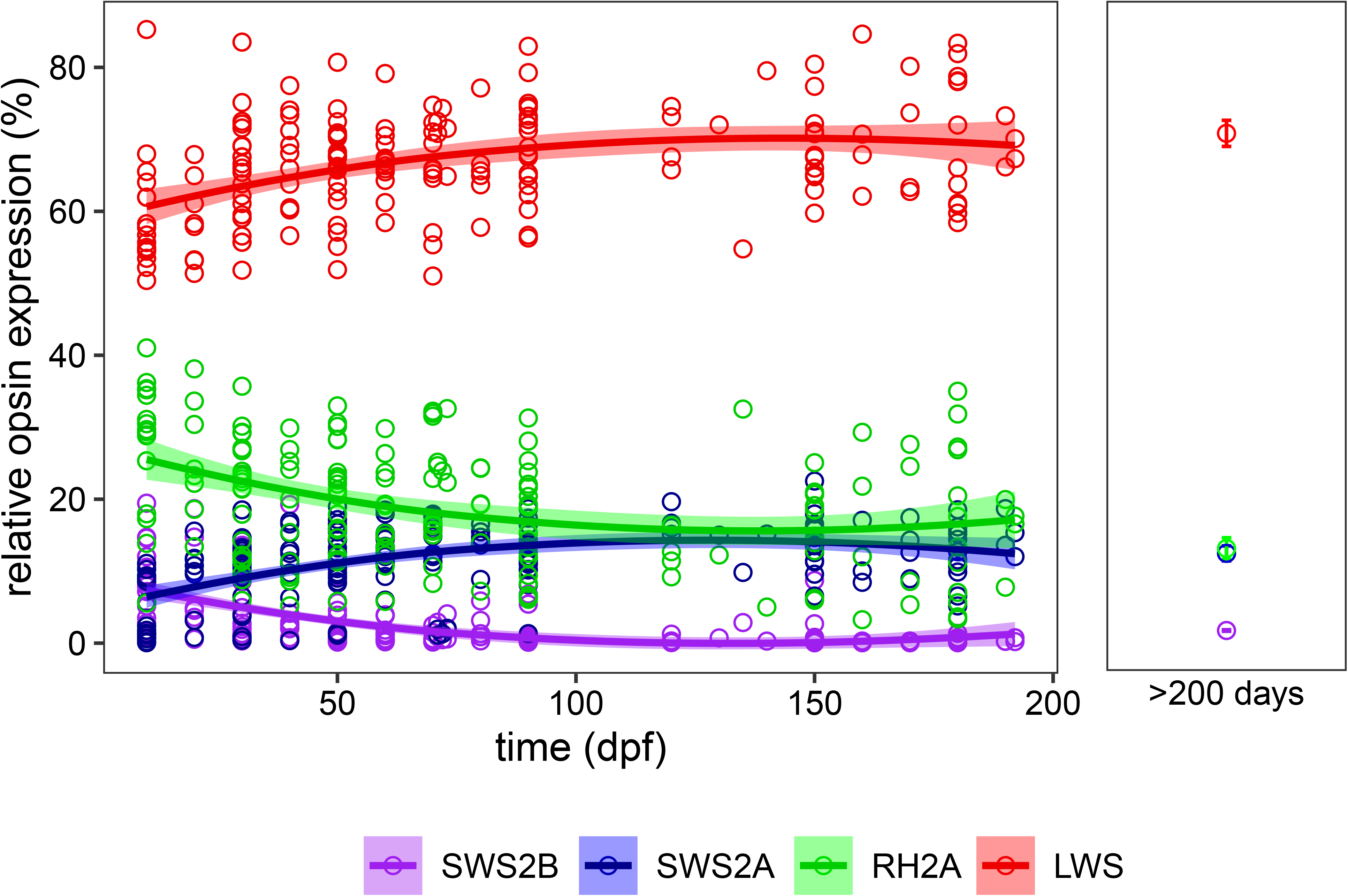
*Pundamilia* opsin expression throughout development. The relative expression of all four opsins (**SWS2B**, **SWS2A**, **RH2A**, and **LWS)** changed during development, as highlighted by the significant effect of time (dpf, days post-fertilization) in all models. Each point represents an individual (2 individuals pooled for time points 10 and 20 dpf), with shaded areas indicating 95% CI. The opsin expression profiles of lab-reared adults (> 200 dpf) are provided for reference (from Wright et al., 2020). Error bars represent the 95% CI.

**Table 1.** Parameter estimates from the models explaining variation in opsin expression. Time represents the days post fertilization, treatment represents the light conditions (broad-spectrum or red-shifted) and species represents the different crosses (*P*.sp. *‘pundamilia-like’* or *P*. sp. *‘nyererei-like’* or their hybrids). We used the *Anova* function (‘car’ package) to estimate the parameter effects, degrees of freedom and P-values of the significant factors in the minimum adequate model. For the removed factors, we used “KRmodcomp” to compare the minimum adequate model with a model containing the removed factor(s).

| **SWS2B** | **F** | **Df** | **Df resid** | **P** |
| --- | --- | --- | --- | --- |
| time | 119.0229 | 1 | 168.23 | < 0.001*** |
| treatment | 91.2334 | 1 | 155.25 | < 0.001*** |
| time:treatment | 5.8824 | 1 | 156.45 | 0.016* |
| **Removed factors** | **F** | **ndf<sup>†</sup>** | **ddf<sup>‡</sup>** | **P** |
| time:species | 0.0396 | 2 | 116.518 | 0.96 |
| treatment:species | 1.1234 | 4 | 30.1961 | 0.36 |
| species | 0.9696 | 2 | 11.2216 | 0.41 |
| **SWS2A** | **F** | **Df** | **Df resid** | **P** |
| time | 14.9456 | 1 | 167.706 | < 0.001*** |
| treatment | 4.7524 | 1 | 153.903 | 0.03* |
| species | 1.3549 | 2 | 10.255 | 0.30 |
| time:treatment | 17.5882 | 1 | 154.815 | < 0.001*** |
| time:species | 2.8392 | 2 | 157.995 | 0.06• |
| **Removed factors** | **F** | **ndf** | **ddf** | **P** |
| treatment:species | 1.8521 | 2 | 152.1398 | 0.16 |

| **RH2A** | **F** | **Df** | **Df resid** | **P** |
| --- | --- | --- | --- | --- |
| time | 21.7602 | 1 | 170.707 | < 0.001*** |
| treatment | 9.5157 | 1 | 157.131 | < 0.01** |
| species | 41.3711 | 2 | 10.154 | < 0.001*** |
| treatment:species | 4.8666 | 2 | 157.615 | < 0.01** |
| **Removed factors** | **F** | **ndf** | **ddf** | **P** |
| time:species | 0.6172 | 2 | 156.1669 | 0.54 |
| time:treatment | 2.0496 | 1 | 156.2806 | 0.15 |
| **LWS** | **F** | **Df** | **Df resid** | **P** |
| time | 30.2459 | 1 | 167.906 | < 0.001*** |
| treatment | 0.2872 | 1 | 156.235 | 0.59 |
| species | 25.8555 | 2 | 9.853 | < 0.001*** |
| time:treatment | 4.0317 | 1 | 156.362 | < 0.05* |
| treatment:species | 3.2048 | 2 | 156.597 | < 0.05* |
| **Removed factors** | **F** | **ndf** | **ddf** | **P** |
| time:species | 0.3604 | 2 | 153.7226 | 0.7 |

† Numerator degrees of freedom
‡ Denominator degrees of freedom

### Species differences

Expression levels of LWS and RH2A differed significantly between the species groups (Table 1, Fig. 2), and post hoc tests showed that all pairwise comparisons were significant (all P < 0.001). LWS expression was highest in *P*. sp. *‘pundamilia-like’*, lowest in *P*. sp. *‘nyererei-like’* and the hybrids were intermediate. RH2A showed the opposite pattern: it was highest in *P*. sp. *‘nyererei-like’*, lowest in *P*. sp. *‘pundamilia-like’* and the hybrids were in-between. Expression levels of the short wavelength sensitive opsins were similar for both species and their hybrids (Table 1).

**Figure 2.**
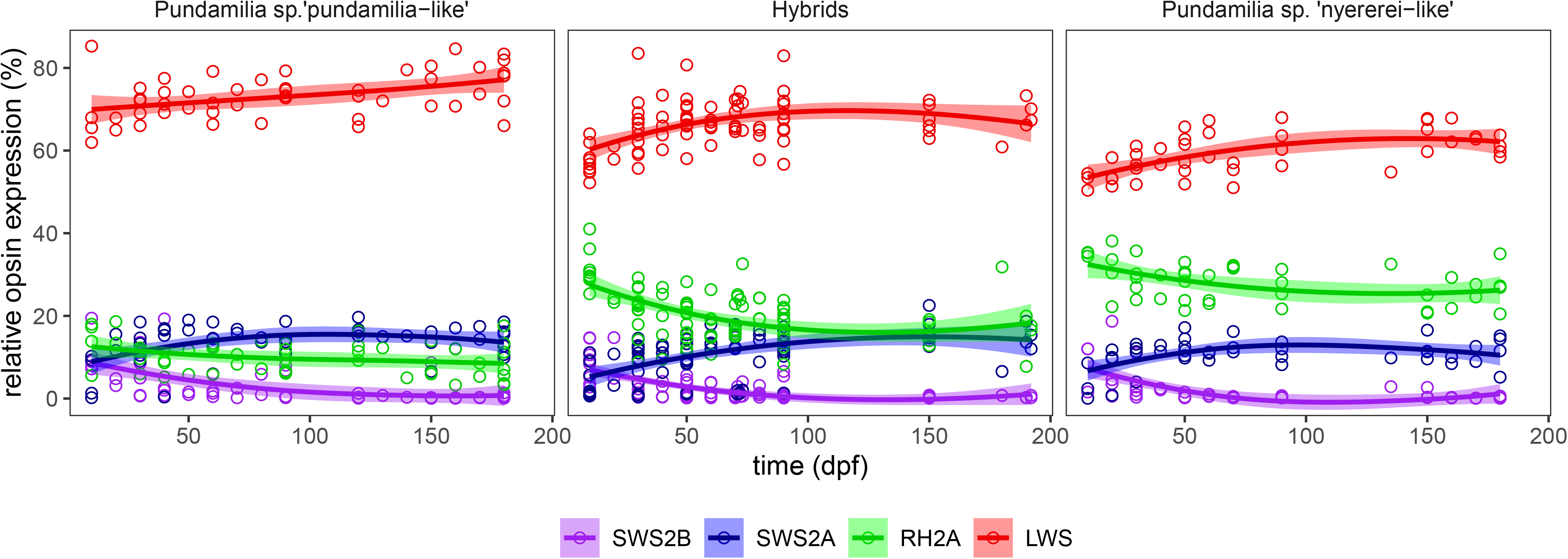
Opsin expression throughout development for *P*. sp. *‘pundamilia-like’*,*P*. sp. *‘nyererei-like’* and their hybrids. LWS and RH2A expression differed significantly between the species groups, but the developmental patterns were similar.

Despite the differences in relative opsin expression levels, there were no differences in the developmental patterns between species: the interaction between species and time did not significantly affect the expression level of any of the opsins (Table 1; there was a trend for SWS2A).

### Effects of the light environment

We found significant effects of the light treatments on SWS2B, SWS2A and RH2A (see Fig. 3 and Table 1). In broad-spectrum light, fish expressed more SWS2B and less SWS2A and RH2A compared to their counterparts from the red-shifted light environment.

**Figure 3.**
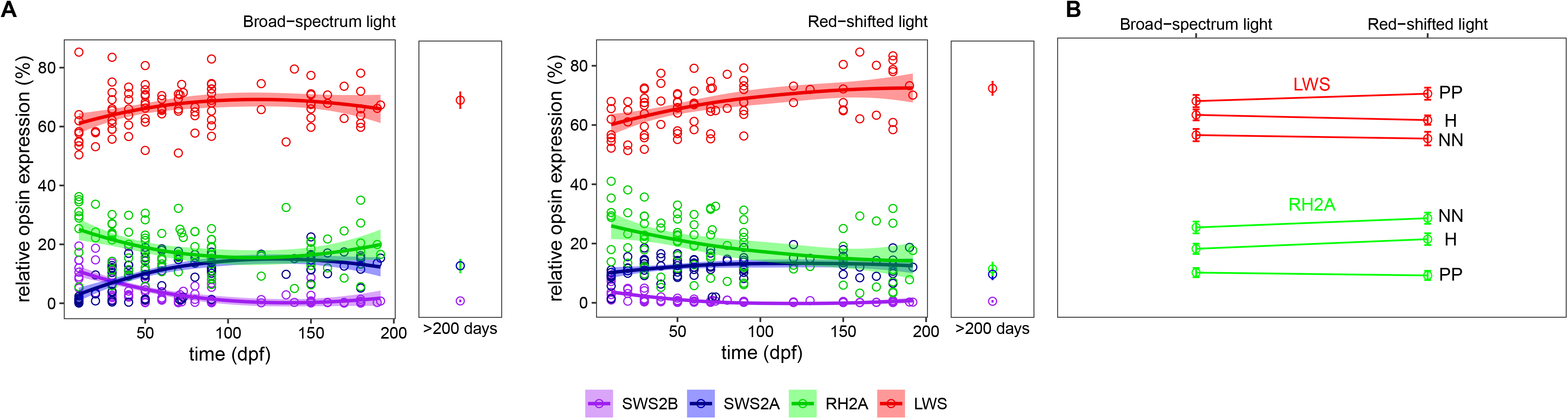
Effects of the light treatments on the developmental patterns of *Pundamilia* opsin expression. (A) The opsin expression patterns of fish housed in broad-spectrum (left panel) vs. red-shifted light (right panel). The expression of SWS2B, SWS2A and RH2A were significantly affected by the light treatments. The developmental patterns of SWS2B, SWS2A and LWS differed significantly between light environments. Shaded areas indicate 95% CI. The opsin expression profiles of lab-reared adults (> 200 dpf) is provided for reference (from Wright et al., 2020). (B) There were significantly different effects of the light treatments on RH2A and LWS expression in *P*. sp. *‘pundamilia-like’*, *P*. sp. *‘nyererei-like’* and their hybrids. Error bars represent the 95% CI.

The developmental patterns of opsin expression were similar between light conditions (Fig. 3), but the rate of change in opsin expression differed significantly. Overall, SWS2B decreased with age in both light conditions, but in broad-spectrum light, SWS2B expression was initially higher and decreased faster than in the red-shifted light condition (Fig. 3). SWS2A, on the other hand, increased in both light conditions, but the rate of change also differed (Table 1). In broad-spectrum light, SWS2A expression was initially lower, but it rapidly increased (during 10 - 50 dpf) until the expression levels were similar in both light conditions. The increase in SWS2A expression was less steep in the red-shifted light condition, where expression remained relatively stable throughout development. LWS expression was lower in early development in the red-shifted light condition and increased more steeply than in the broad-spectrum light environment. The developmental pattern of RH2A expression was not affected by the light treatments.

We also examined whether the two *Pundamilia* species and their hybrids responded differently to the light treatments. We found significant interaction effects (species by treatment) for RH2A and LWS expression (Table 1, Fig. 3). *P*. sp. *‘nyererei-like’* and the hybrids had higher RH2A expression in the red-shifted light environment, while *P*. sp. *‘pundamilia -like’* RH2A expression did not differ between conditions. LWS showed the opposite pattern: in *P*. sp. *‘nyererei-like’* and the hybrids it did not differ between light conditions, but *P*. sp. *‘pundamilia-like’* had higher LWS expression in the red-shifted light environment.

### Hybrids

We also explored whether the reciprocal hybrids differed in opsin expression (Table 2, Fig. 4). For this analysis, we used samples ≤ 90 dpf, as we lacked older samples for the NP hybrids (NP: *P*. sp. *‘nyererei-like’* mother, *P*. sp. *‘pundamilia-like’* father; PN*: P*. sp. *‘pundamilia-like’* mother, *P*. sp. *‘nyererei-like’* father). Figure 4 suggests that hybrids tend to resemble their maternal species more than their paternal species, particularly in SWS2A and RH2A expression.

**Table 2.** Parameter estimates from the models explaining variation in opsin expression among the reciprocal hybrids until 90 dpf. Time represents days post fertilization, treatment represents the light conditions (broad-spectrum or red-shifted) and cross denotes NP (*P*. sp. *‘nyererei-like’* mother, *P*. sp. *‘pundamilia-like’* father) or PN (*P*. sp. *‘pundamilia-like’* mother, *P*. sp. *‘nyererei-like’* father). We used the *Anova* function to estimate the parameter effects, degrees of freedom and the P-values of the significant factors in the minimum adequate model. For the removed factors, we used a “KRmodcomp”.

| **SWS2B** | **F** | **Df** | **Df resid** | **P** |
| --- | --- | --- | --- | --- |
| time | 38.687 | 1 | 64.318 | < 0.001 *** |
| treatment | 52.458 | 1 | 60.929 | < 0.001 *** |
| **Removed factors** | **F** | **ndf<sup>†</sup>** | **ddf<sup>‡</sup>** | **P** |
| time:treatment | 0.0905 | 1 | 60.1676 | 0.76 |
| treatment:cross | 0.5921 | 2 | 9.81 | 0.57 |
| time:cross | 2.0146 | 1 | 58.2417 | 0.16 |
| cross | 0.0812 | 1 | 4.1869 | 0.79 |
| **SWS2A** | **F** | **Df** | **Df resid** | **P** |
| time | 13.5463 | 1 | 63.36 | < 0.001 *** |
| treatment | 3.2938 | 1 | 61.152 | 0.07• |
| cross | 4.8588 | 1 | 3.549 | 0.10 |
| time:treatment | 9.81 | 1 | 61.328 | < 0.01 ** |
| **Removed factors** | **F** | **ndf** | **ddf** | **P** |
| time:cross | 0.1285 | 1 | 63.4182 | 0.72 |
| treatment:cross | 1.7816 | 1 | 62.5275 | 0.19 |
| **RH2A** | **F** | **Df** | **Df resid** | **P** |
| time | 15.1876 | 1 | 65.191 | < 0.001 *** |
| treatment | 8.5974 | 1 | 61.882 | < 0.01 ** |
| cross | 1.7027 | 1 | 3.696 | 0.27 |
| time:cross | 4.7041 | 1 | 65.345 | 0.03 * |
| **Removed factors** | **F** | **ndf** | **ddf** | **P** |
| time:treatment | 0.0002 | 1 | 60.5536 | 0.99 |
| treatment:cross | 0.6981 | 1 | 63.9932 | 0.41 |
| **LWS** | **F** | **Df** | **Df resid** | **P** |
| time | 11.915 | 1 | 68.32 | < 0.001 *** |
| **Removed factors** | **F** | **ndf** | **ddf** | **P** |
| treatment:cross | 1.4291 | 3 | 15.0737 | 0.27 |
| time:treatment | 1.3407 | 1.3407 | 63.332 | 0.25 |
| treatment | 3.4179 | 1 | 63.2963 | 0.07• |
| time:cross | 2.6334 | 1 | 25.3959 | 0.12 |
| cross | 0.0664 | 1 | 3.7826 | 0.81 |

<sup>†</sup> Numerator degrees of freedom
<sup>‡</sup> Denominator degrees of freedom

**Figure 4.**
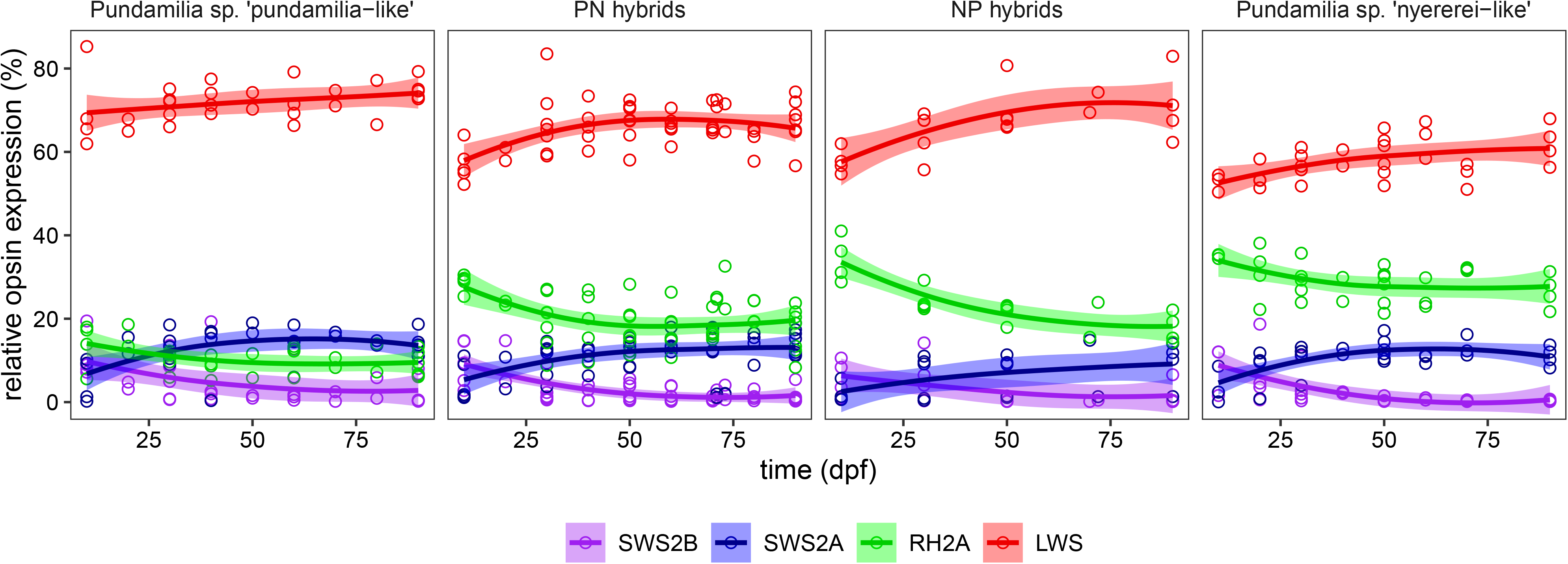
Differences in the developmental patterns of opsin expression between reciprocal hybrids. RH2A was the only opsin that differed across time for the reciprocal hybrids; the NP hybrids had a faster decrease in their expression. The *P*. sp. *‘pundamilia-like’* and *P*. sp. *‘nyererei-like’* panels show that the PN hybrids resemble *P*. sp. *‘pundamilia-like’* and the NP hybrids resemble *P*. sp. *‘nyererei-like’*. Shaded areas indicate 95% CI.

This effect was statistically significant only for RH2A, showing an interaction of time and cross type, indicating a difference in developmental pattern between PN and NP hybrids. Both hybrid types had similar RH2A expression at 90 dpf, but the NP hybrids started with high expression and rapidly decreased, similar to *P*. sp. *‘nyererei-like’*. The PN hybrids exhibited a more stable and lower expression level, similar to *P*. sp. *‘pundamilia-like’*.

## Discussion

In this study, we characterised the developmental pattern of opsin expression in two Lake Victoria haplochromines, *P*. sp. *‘pundamilia-like’* and *P*. sp. *‘nyererei-like’*, and their hybrids in two distinct light conditions. Our results show that opsin expression changes over developmental time and is also influenced by environmental light.

### *Pundamilia* opsin expression throughout development

We hypothesised that the relative opsin expression in *Pundamilia* cichlids (SWS2B, SWS2A, RH2A and LWS) would remain fairly constant across development, with relatively high long-wavelength sensitivity throughout, as previously documented in sand-dwelling cichlids from Lake Malawi (Carleton et al., 2008). Our results are partly consistent with this hypothesis: throughout development, expression of the long-wavelength sensitive opsin (LWS) was high, compared to the medium- and short-wavelength sensitive opsins (RH2A, SWS2B and SWS2A). However, contrary to the Lake Malawi sand-dwellers, opsin expression in *Pundamilia* changed over development. This resembles the developmental pattern of *O. niloticus* (Carleton et al., 2008; O’Quin et al., 2011). Therefore, the developmental pattern of *Pundamilia* cichlids falls in-between *O. niloticus* and Lake Malawi sand-dwellers. When comparing our present findings to the opsin expression levels we documented in adults (Wright et al., 2020), we observe that the opsin expression profile stabilizes around 200 dpf.

At 10 dpf, SWS2B expression was relatively high but quickly decreased to zero at ~100 dpf. A possible adaptive explanation for high SWS2B expression in young fish might be an ontogenetic shift in the vertical distribution in the water column. In several haplochromine species from Lake Victoria, larvae and juveniles have shallower depth distributions than adults (Goldschmidt, Witte, & De Visser, 1990) and shallow waters are relatively rich in short-wavelength light (captured by SWS1, SWS2B and SWS2A). The depth distribution of *Pundamilia* juveniles is unknown, but high SWS2B expression may suggest that they inhabit relatively shallow waters. SWS2B expression may also contribute to foraging efficiency; cichlids and other teleost fish change their foraging strategies as they develop, and zooplankton is an important component of larval and juvenile diets (Fryer & Iles, 1972). Short-wavelength vision particularly aids the detection of small and translucent objects (Britt, Loew, & McFarland, 2001; Carleton et al., 2008; Loew & Wahl, 1991; Novales-Flamarique & Hawryshyn, 1994; Novales Flamarique, 2013).

### Species differences

The parental species differed in LWS and RH2A expression: *P*. sp. *‘pundamilia-like’* had higher LWS expression than *P*. sp. *‘nyererei-like’*, while RH2A exhibited the opposite pattern. This is in line with earlier studies of wild caught (Wright et al., 2019) and laboratory-reared fish (Hofmann et al., 2009, Wright et al., 2020). As hypothesised, the developmental patterns of opsin expression were similar in both species.

### Effects of the light environment

The fish were reared in two light environments mimicking the natural shallow (broad-spectrum) and deep (red-shifted) light environments of Lake Victoria. These light treatments influenced opsin expression. SWS2B expression was higher in broad-spectrum light than in red-shifted light, while SWS2A and RH2A followed the opposite pattern. Higher SWS2A expression in the red-shifted condition contrasts with what was previously observed in *Pundamilia* adults (Wright et al., 2020); we discuss this further below.

In addition to the overall expression levels, the developmental patterns of LWS, SWS2A, and SWS2B were affected by our light treatments. LWS expression increased faster in the red-shifted light condition, similar to the pattern observed in Midas cichlids (Härer et al., 2017). However, the SWS opsins showed a different effect, changing faster in the broad-spectrum light conditions: SWS2A increased faster and SWS2B decreased faster. Thus, our findings do not correspond to the general pattern suggested for neotropical cichlids, where progress towards the adult phenotype was accelerated in light conditions rich in long wavelengths (Härer et al., 2017). Rather, it seems that LWS and SWS2A increase faster in the light conditions where these opsins confer greater photon capture. To test this interpretation, experiments are needed to quantify fish’ visual performance in different visual conditions and at different developmental stages.

When analysing the interaction between environmental light and species (*P*. sp. *‘pundamilia-like’*, *P*. sp. *‘nyererei-like’* or hybrids), we found that LWS and RH2A expression were both species- and environment-specific (Fig. 3). *P*. sp. *‘pundamilia-like’* reared in the red-shifted light environment expressed more LWS than their counterparts raised in broad-spectrum light. *P*. sp. *‘nyererei-like’* and hybrid LWS expression did not differ between the treatments. RH2A showed the opposite expression pattern between species: *P*. sp. *‘nyererei-like’* and the hybrids had higher RH2A expression in red-shifted light, while *P*. sp. *‘pundamilia-like’* expression did not differ between the treatments. These results suggest that both *Pundamilia* species respond plastically to the light treatments, but in different ways. In the adults, LWS and RH2A were also differently expressed between species in the two light environments (Wright et al., 2020). However, contrary to what we observed here, adult *P*. sp. ‘*nyererei-like’* expressed more LWS in the red-shifted condition than in the broad-spectrum condition. Hybrids only differed in SWS2A expression and *P*. sp. ‘*pundamilia-like’* did not show any significant differences. The mismatch between our results and the data from the adults might be explained by the lack of data points between 180 dpf and the adult stage (200 dpf and older). Possibly, *P*. sp. *‘nyererei-like’* increase their LWS expression during this time. Additional sampling during this developmental period, as well as transfer experiments (e.g.(Härer, Karagic, Meyer, & Torres-Dowdall, 2019)), are required to establish the sensitive window for the effects of light conditions on opsin expression and potential species differences in this regard.

Heterochronic changes are modifications in developmental pattern compared to the ancestral state (McKinney & McNamara, 1991). For haplochromine cichlids, *O. niloticus* is considered to represent the ancestral pattern of opsin expression development (Carleton et al., 2008; O’Quin et al., 2011). Rock- and sand-dwelling species from Lake Malawi differ greatly from *O. niloticus*, and differences in their developmental patterns can be interpreted as adaptations to different light environments. *P*. sp. *‘pundamilia-like’* and *P*. sp. *‘nyererei-like’* also differ from *O. niloticus*. Yet, even though the two *Pundamilia* species inhabit very different natural light environments, there is little difference in their developmental patterns. We do find that they respond somewhat differently to the light treatments. This may represent a starting point for heterochronic shifts in *Pundamilia*, possibly providing a target for selection and allowing the evolution of species-specific developmental trajectories.

### Hybrids

The developmental profiles of hybrids fell in-between the parental species. The level of environmental plasticity in opsin expression was similar between hybrids and parental species, in contrast to our prediction that hybrids might show greater plasticity, but in line with the earlier data on adults (Wright et al., 2020). We used the reciprocal hybrids (≤ 90 dpf only) to test for potential parental effects on opsin expression. Our results suggest that PN (*P*. sp. *‘pundamilia-like’* mothers) and NP hybrids (*P*. sp. *‘nyererei-like’* mothers) may have different developmental profiles: RH2A expression decreased faster in NP hybrids than in PN hybrids. This might be interpreted as a parental effect, as PN hybrids showed more similar expression patterns to *P*. sp. *‘pundamilia-like’* (higher SWS2A and lower RH2A), while NP hybrids showed more similar patterns to *P*. sp. *‘nyererei-like’*. Thus, while our study was not designed to fully explore the influences of maternal or paternal effects on opsin expression, it provides some exciting first indications that warrant further investigation.

## Conclusion

The evolutionary importance of heterochrony lies in the fact that certain phenotypes appear during development and are beneficial in a given environment, which can then become targets of selection, potentially leading to arrested or accelerated development. Likewise, environmentally-induced phenotypes, throughout development, are ‘seen’ by natural selection and can provide stepping stones for evolutionary adaptation (West-Eberhard, 2003).

In this study, we documented the developmental pattern of opsin expression in two species of Lake Victoria cichlids and their reciprocal hybrids.

We found that *Pundamilia* cichlids progress from high levels of short-wavelength-sensitive opsin expression as larvae and juveniles to high levels of long-wavelength-sensitive opsin expression as (sub)adults. This pattern may reflect differences between life stages in water depth distribution, where larvae and juveniles reside in more shallow waters, with broad-spectrum light, compared to adults. Developmental patterns were similar between the species, but the overall opsin expression levels differed between them, consistent with prior work. The developmental trajectory of opsin expression also responded plastically to the visual conditions. Finally, we found preliminary evidence for parental effects in the developmental patterns of opsin expression that warrants additional research.

## Acknowledgements

We acknowledge Sjoerd Veenstra and Brendan Verbeek for taking care of the fish. Financial support came from the Swiss National Science Foundation (SNSF PZ00P3-126340; to MM), the Netherlands Foundation for Scientific Research (NWO VENI 863.09.005; to MM) and the University of Groningen.

