## Supplemental Tables 1, 2 and 3. Supplemental Figure 1 for "Developmental and environmental plasticity in opsin gene expression in Lake Victoria cichlid fish"

Supplementary information

**Table S1. F1 and F2 families used for the opsin expression analysis.** Samples from each cross reared in the broad-spectrum light condition (mimicking shallow water; ‘S’) and the red-shifted light condition (mimicking deep water; ‘D’). The family name is indicated as mother and as father, such that PN indicates a F1 family from *P.* sp. *‘pundamilia-like’* mother and *P.* sp. *‘nyererei-like’* father. Four letters (PPPP) indicate a F2 family (i.e., a PP mother crossed with a PP father). The reciprocal crosses (PN and NP) are grouped as hybrids. Superscript numbers indicate families with the same mother, and superscript letters indicate families with the same father.

| ***P.* sp. *‘pundamilia-like’*** | | | **Hybrids** | | | | | | ***P.* sp*. ‘nyererei-like’*** | | |
| --- | --- | --- | --- | --- | --- | --- | --- | --- | --- | --- | --- |
| **family** | **S** | **D** | **family** | **S** | **D** | **family** | **S** | **D** | **family** | **S** | **D** |
| PP16 | 1 | 1 | PN11^1^ | 4 | 3 | NP8^5e^ | 1 | 0 | NNNN1 | 6 | 6 |
| PP17^1a^ | 4 | 3 | PN12^1^ | 5 | 4 | NP9^5e^ | 2 | 0 | NNNN2^6g^ | 5 | 6 |
| PP18^1a^ | 5 | 7 | PPNN2^3d^ | 3 | 5 | NNPP1^f^ | 5 | 5 | NNNN3^g^ | 4 | 5 |
| PPPP5^2^ | 1 | 3 | PPNN3^d^ | 4 | 4 | NNPP2 | 5 | 5 | NNNN4 | 5 | 5 |
| PPPP6^b^ | 3 | 3 | PPNN4^d^ | 2 | 4 | NNPP3^f^ | 5 | 4 | NNNN5 | 3 | 1 |
| PPPP7^b^ | 6 | 5 | PPNN5^3^ | 3 | 3 |  |  |  | NNNN6 | 1 | 1 |
| PPPP8^2b^ | 5 | 5 | PPNN6^d^ | 2 | 1 |  |  |  | NNNN7^6^ | 5 | 5 |
| PPPP9 | 1 | 0 | PPNN7^d^ | 3 | 3 |  |  |  |  |  |  |
| PPPP10^c^ | 5 | 4 | PPNN8^4d^ | 3 | 4 |  |  |  |  |  |  |
| PPPP11^c^ | 3 | 4 | PPNN9 | 5 | 5 |  |  |  |  |  |  |
|  |  |  | PPNN10^4d^ | 5 | 5 |  |  |  |  |  |  |
| *total* | 34 | 35 | *total* | 39 | 41 | *total* | 18 | 14 | *total* | 29 | 29 |

| **Dpf** | **#fish** | **#eyes** | **Tissue** |
| --- | --- | --- | --- |
| **10** | 2 | 4 | Whole eye |
| **20** | 2 | 4 | Whole eye |
| **30** | 1 | 2 | Whole eye |
| **40** | 1 | 2 | Whole eye |
| **50** | 1 | 2 | Whole eye |
| **60** | 1 | 2 | Whole eye |
| **70** | 1 | 1 | Whole eye |
| **80** | 1 | 1 | Whole eye |
| **90** | 1 | 1 | Whole eye |
| **120** | 1 | 1 | Retina |
| **140** | 1 | 1 | Retina |
| **150** | 1 | 1 | Retina |
| **160** | 1 | 1 | Retina |
| **180** | 1 | 1 | Retina |

**Table S2. Sampling time points.** Dpf represents the days post fertilization, #fish represents the number of fish sampled and ‘#eyes’ is the number of eye samples used for the RNA isolations of the respective tissues (whole eye or retina).

**Table S3.** **Sequences of the primers and probes used for the qPCR reactions.**

| SWS2B | Forward Primer | 5’-GCGCTGCACTTCCACCTC-3’ |
| --- | --- | --- |
|  | Reverse Primer | 5’-GGCCACAGGAACACTGCAT-3’ |
|  | Probe | 5’-FAM-TTGGATGGAGCAGGTATATCCCAGAGGG-Tamra-3’ |
| SWS2A | Forward Primer | 5’-CAAGATTGAAGGTTTCATGGTA-3’ |
|  | Reverse Primer | 5’-CGCTCGAAAGCTATCACAGC-3’ |
|  | Probe | 5’-FAM-ACTCGGTGGTATGGTAAGCCTGTGG-Tamra-3’ |
| RH2A | Forward Primer | 5’-TTCTGTGCWATTGAGGGATTC-3’ |
|  | Reverse Primer | 5’-CCAGGACAACAAGTGACCAGAG-3’ |
|  | Probe | 5’-FAM-TGGCCACACTWGGAGGTGAAGTTGC-Tamra-3’ |
| LWS | Forward Primer | 5’-CTGTGCTACCTTGCTGTGTGG-3’ |
|  | Reverse Primer | 5’-GCCTTCTGGGTTGACTCTGACT-3’ |
|  | Probe | 5’-FAM-TGGCCATCCGTGCTGTTGC.Tamra-3’ |

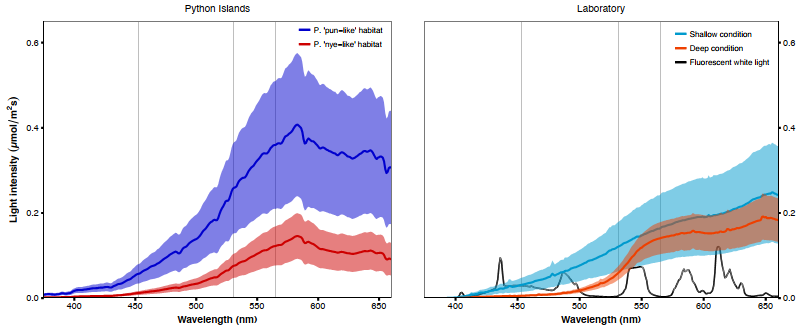

**Figure S1. Light conditions at Python Islands and in the laboratory.** Left panel: Downwelling irradiance in the natural habitats of P. sp. 'pundamilia-like' (0.5-2m depth; blue curve) and P. sp. 'nyererei-like' (0.5-5m depth, red curve). Right panel: Downwelling irradiance in the ‘shallow’ (blue curve) and ‘deep’ (red curve) light treatments in the laboratory. Curves represent averages of multiple measurement series with standard errors. Grey vertical lines indicate the maximum sensitivity of the three main photoreceptors of Pundamilia: LWS (565nm), RH2 (531 nm) and SWS2A (453nm) (based on (Carleton, Parry, Bowmaker, Hunt, & Seehausen, 2005; Wright et al., 2017)
